# AI Derived Therapeutic Development for the Treatment of Opioid Use Disorder

**DOI:** 10.1101/2024.02.26.579391

**Authors:** Valeria Lallai, A.C. Martin, J.P. Fowler, Malia Bautista, Allison S. Mogul, Jinjutha E. Cheepluesak, Saman Mirzaei, Ian Jenkins, Jonathan R.T. Lakey, Robert Tinder, Christie D. Fowler

## Abstract

The opioid epidemic has led to a devastating loss of life nationwide. Of those dependent on opioids, many individuals desire to quit or reduce use, but their efforts are often unsuccessful given the powerful reinforcing properties associated with opioid drugs, especially fentanyl given its high potency and speed of onset. Here, we developed a novel theraputic based on a newly developed artificial intelligence (AI)-based platform, which was rationally designed to identify markers of dysregulation from human drug user postmortem brain tissue. The GATC-021 compound was synthesized and validated with *in vitro* screening for target specificity. Thereafter, GATC-021 was examined for its effectiveness in modulating opioid dependence with an animal model of addiction. We found that GATC-021 substantially reduced fentanyl intake in both male and female rats, as assessed with intravenous self-administration. However, given drug soluability challenges, additional studies are needed to better develop drug formulations to permit translation into clinical populations more effectively. Taken together, these findings validate our AI-based platform for novel therapeutic development with a polypharmacy approach and further support the effectiveness of such target modulation as a promising therapeutic approach for those suffering from opioid use disorder.

## INTRODUCTION

Dysregulated opioid use has led to devastating consequences for the lives of many individuals, affecting millions worldwide^1^. In 2017, the opioid epidemic was declared a national emergency in the US^2^. Approximately three million individuals in the USA currently suffer from opioid use disorder (OUD)^3^. Alarmingly, the vast majority of recent deaths have been attributed to synthetic opioids such as fentanyl. Current treatment options for OUD include methadone, buprenorphine, naloxone^1^. However, these medications are significantly limited by suboptimal safety margins, reduced long-term efficacy, and/or treatment adherence challenges. Thus, the significant morbidity, mortality, and economic consequences imposed by opioid misuse represents a pressing global concern that calls for novel and innovative treatment strategies.

Fentanyl is a synthetic analgesic that exhibits high potency and lipophilic properties, leading to relatively quick pharmacokinetic actions in the brain and thereby inferring fast analgesic properties, high potential for respiratory depression, and high addiction liability^3,4^. Fentanyl exhibits agonist properties with direct binding to the μ-, κ-, and 8-opioid receptors, leading to downstream intracellular changes in β arrestin and G-protein signaling cascades^4–7^. Recent *in vitro* evidence also suggests that fentanyl may act on the α1 adrenoceptor subtypes, dopamine D1 and D4 receptors, and serotonin 5-HT1A and 5-HT2A receptors, in addition to blocking catecholamine uptake through actions on the vesicular monoamine transporter 1^5,8–10^. Interestingly, with serotonergic signaling, fentanyl has been shown to modulate activity of a recently identified μ-opioid/5-HT1A receptor heterodimer at the plasma membrane *in vitro*, in which fentanyl increased the downstream intracellular 5-HT1A signaling pathways for MAPK p38 and Erk1/2^11^. Moreover, long-term treatment with fentanyl appears to induce persistent changes in receptor and related circuit function, for instance as evidenced at the μ-opioid receptor level with tolerance and increased desensitization^12^. The development of clinical symptoms related to chronic opioid abuse, including dependence, tolerance and addiction, are thought to involve opioid receptors within the mesolimbic reward system^13^. Specifically, μ-and δ-opioid receptors agonists, such as fentanyl, increase dopaine release from ventral tegmental area (VTA) neurons projecting into the nucleus accumbens (NAc), which is thought to underlie the reinforcing properties of the drug^14^, whereas tolerance and withdrawal-mediated effects are thought to involve inhibitory signaling within and into the VTA^15–17^. Importantly, 5-HT receptors are also localized within addiction-associated brain regions, including the VTA, NAc and prefrontal cortex^18,19^. Therefore, it is of particular interest to target pathways involved in the molecular changes associated with opioid’s effects as an approach for the development of more effacacious therapeutics.

Artificial or augmented intelligence (AI) therapeutic strategies have rapidly emerged across the healthcare market and currently represent a source of innovation within various fields, including neurology, oncology, and cardiology. Here, we employed the GATC multi-omics platform, which was developed using a complex integration of large genomic, transcriptomic, proteomic and metabolomic datasets, that were generated from postmortem human brain tissue in patients dependent on opioids. From the data, 20 high probability causative biomarkers were identified by Liquid Bioscience and subsequently delivered to GATC Health platform to analyze the systems biology and predict the treatment markers and mechanisms. Using the GATC AI platform we identified a polypharmacological target profile in which we sought to validate utilizing known literature compounds. Specifically, based on our AI output we predicted that a compound comprising of agonist activity on the serotonin receptors 5-HT2A and 5-HT6, along with increased thiamine, would be effacacious in mitigating drug taking behavior. This analysis led to the identification of a family of >80 lead compounds, which were rationally identified as targeting processes dysregulated in OUD, and GATC-021 was selected as a starting point for these studies given its corresponding classification as a non-hallucinogenic psychedelic analogue in a prior report (e.g., compound 14 in Cameron et al^20^). We thus first examined the effectiveness of GATC-021 to modulate synaptic targets predicted to mitigate the long-term effects of opioid use with *in vitro* assays. Thereafter, the GATC-021 compound along with the prodrug sulbutiamine were examined for their effectiveness in mitigating fentanyl intake with intravenous self-administration, the ‘gold-standard’ rodent model of drug taking behavior with high translational relevance to human patterns of drug use. Further, control studies were performed to assess whether GATC-021 combined with sulbutiamine affected overall behavioral measures and/or induced toxcity. Taken together, our studies validate the AI platform for the identification of novel targets for therapeutic development and further provide evidence that AI directed methods can serve as a promising tools for novel therapeutic design and biological targeting of OUD.

## METHODS

### Synthesis and Preparation of GAT-021

hydrochloride, were purchased from Ambeed. 37% hydrochloric acid was obtained from Stellar Chemicals. Sulfuric acid, dichloromethane and palladium catalyst was purchased from Sigma-Aldrich. Ethanol (denatured with 5% isopropanol) and sodium hydroxide were obtained from Oakwood Chemicals. Methanol, isopropanol, and reagents were obtained from VWR. Column chromatography was performed on an Isco Combiflash instrument using columns as described. ^1^H and ^13^C NMR spectra were recorded on either Varian or JEOL 400 MHz NMR spectrometers. NMR analysis was performed in deuterated solvents purchased from Cambridge Isotope Laboratories. Chemical shifts (δ) are reported in parts per million and referenced to the internal standard, tetramethylsilane (TMS), and/or the residual solvent peaks (e.g., CHCl_3_). Liquid Chromatography-Mass Spectrometry (LC-MS) data were obtained on either a Shimadzu 2020, Nexera X2 or, a Waters Acquity UPLC equipped with an SQD detector. High-resolution mass spectrometry (HRMS) data were obtained on a Hybrid Quadrupole-Orbitrap Mass Spectrometer or APCI-Q-TOF (**Supplementary Material**).

Synthesis of 9-(benzyloxy)-3-methyl-1,2,3,4,5,6-hexahydroazepino[4,5-*b*]indole: without further purification or characterization.

Synthesis of 3-methyl-1,2,3,4,5,6-hexahydroazepino[4,5-*b*]indol-9-ol: hexahydroazepino[4,5-*b*]indol-9-ol, (81.7%) as a dark colored oil.

^1^H NMR (400 MHz, DMSO-*d*) δ 10.31 (s, 1H), 6.99 (d, *J* = 8.5 Hz, 1H), 6.64 (d, *J* = 2.3 Hz, 1H), 6.47 (dd, *J* = 8.5, 2.3 Hz, 1H), 6.02 (s, 1H), 2.84 – 2.80 (m, 2H), 2.67 (s, 6H), 2.38 (s, 3H).

^13^C NMR (126 MHz, DMSO-*d*) δ 150.11, 137.51, 129.18, 128.88, 110.67, 110.29, 109.75, 101.39, 57.71, 55.91, 48.59, 45.51, 27.71, 23.58.

HRMS (ESI +) calculated for C_13_H_16_N_2_O^+^ [M + H^+^] 217.1335; found 217.1332 LCMS m/z: 217.10 [M+H]+

Preparation of 3-methyl-1,2,3,4,5,6-hexahydroazepino[4,5-*b*]indol-9-ol hydrochloride:

Synthesis of 3-methyl-1,2,3,4,5,6-hexahydroazepino[4,5-*b*]indol-9-ol hydrochloride, (GATC-021): 12.1 grams of 3-methyl-1,2,3,4,5,6-hexahydroazepino[4,5-*b*]indol-9-ol was redissolved in 150 mL of 40°C isopropanol, when 6.75 mL of 37% aqueous HCl was added. The mixture rapidly formed a black precipitate which upon cooling to room temperature was then filtered off, to afford 14.0 grams of 3-methyl-1,2,3,4,5,6-hexahydroazepino[4,5-*b*]indol-9-ol hydrochloride, (99.1%) after drying overnight under high vacuum.

### In Vitro Studies

Assays were performed at Eurofins DiscoverX (Fremont, Calif., USA). GATC-021 was examined with the following: 5-HT2A Human Serotonin GPCR Cell Based Agonist Arrestin Assay (Cat # 86-0001P-2090AG), 5-HT2B Human Serotonin GPCR Cell Based Agonist Calcium Flux Assay (Cat # 86-0030P-2091AG), 5-HT6 Human Serotonin GPCR Cell Based Agonist Arrestin Assay (Cat # 86-0001P-2094AG), 5-HT7 Human Serotonin GPCR Cell Based Agonist cAMP Assay (Cat # 86-0007P-2255AG), and TRKB Human RTK Kinase Cell Based Agonist Functional Assay (Cat # 86-0006P-2761AG). For all of the above, the final assay vehicle concentration was 1%, and the results are expressed as a percent efficacy relative to the control ligand.

### *In Vivo* Studies

#### Animals

Adult male and female Wistar rats were purchased from Charles River. The subjects were housed in an environmentally controlled vivarium on a 12-hour reversed light:dark cycle. Food and water were provided ad libitum until behavioral training commenced. All testing was conducted during the dark phase of the light cycle, when rats are most active. Food and water were also provided ad libitum during all procedures. All procedures were conducted in strict accordance with the NIH Guide for the Care and Use of Laboratory Animals and were approved by the Institutional Animal Care and Use Committee of the University of California, Irvine.

#### Drugs

Fentanyl (Cat # F3886, Sigma) was dissolved in 0.9% sterile saline for intravenous self-administration. Sulbutiamine (Cat # S09645G, Fisher Scientific) was dissolved in corn oil and injected subcutaneously at a concentration of 50 mg/ml. The GATC-021 compound was dissolved in a vehicle solution (10% DMSO, 10% Tween-80, and 80% sterile saline), pH was adjusted to ∼7-7.4, and injected intraperitoneally.

#### Open Field Test

To examine the effects of GATC-021 on general behavior, subjects (n=6; 3 male, 3 female) were injected and then placed in a large square arena to freely explore the open field. The chamber was composed of plexiglass (35 cm L x 35 cm W x 31 cm H) as described previously^21,22^, with a shielded white light lamp ∼90 cm above the apparatus for consistent lighting. Prior to testing, animals were habituated by handling for at least 5 mins per day for 2 days. On the test day, they were injected with sulbutiamine (50mg/kg, sc) and then 20 min later, GATC-021 (0, 50 or 70 mg/kg, ip). Twenty min after the GATC-021 injection, rats were then individually placed into the open field and recorded for a 15 min test. Behavior was video recorded and quantified with the unbiased software system, AnyMaze.

#### Fentanyl self-administration

After their arrival, subjects were habituated in the UCI vivarium for 3-5 days. Thereafter, male and female rats were anesthetized with an isoflurane (1%– 3%)/oxygen vapor mixture and prepared with intravenous catheters, as previously described^23^. These catheters consisted of a 14 cm length of silastic tubing fitted to a guide cannula (Plastics One), which was bent at a curved right angle and encased in dental acrylic. The catheter tubing was passed subcutaneously from the animal’s back to the right jugular vein, and a 2 cm length of the catheter tip was inserted into the vein and secured with surgical silk suture. Following the surgery, subjects were provided a post-surgical recovery period of >48 hours. On the first day of self-administration, male and female rats were connected to the tubing port on the back to permit self-administration of fentanyl infusions at a dose of 2.5 μg/kg/infusion (n=44; 22 male, 22 female) or saline (n=12; 6 male, 6 female). The saline self-administration group served as control for fentanyl. Infusions were delivered through the tubing into the intravenous catheter by a Razel syringe pump (Med Associates). Training began at a fixed ratio 1, time out 20 sec (FR1TO20) schedule of reinforcement during 1-hour daily sessions, and once the subjects reached the criteria of 10 infusions in one session, the fixed ratio schedule was increased to achieve a final fixed ratio 2 (FR2TO20) schedule of reinforcement. Operant responding was assessed using two retractable levers (one active, one inactive) that extended into the chamber (MedAssociates). Completion of the response criteria on the active lever resulted in the delivery of a drug infusion. Responses to the inactive lever resulted in no scheduled consequence. Since we were interested in examining the effects of chronic fentanyl intake, subjects were provided access to stabilize their responding for the fentanyl infusions across 10 daily sessions. On Day 11, intraperitoneal injections with the GATC-021 compound were initiated prior to the daily fentanyl self-administration session, and this pairing of daily GATC-021 compound injections followed by intravenous fentanyl self-administration continued across 5 consecutive daily sessions. Doses were chosen based on the results obtained studying the effects of GATC-021 on general behavior. Groups included: (1) Fentanyl self-administration with vehicle (0 mg/kg GATC-021) and sulbutiamine, (2) Fentanly self-administration with 25 mg/kg GATC-021 and sulbutiamine, (3) Fentanyl self-administration with 50 mg/kg GATC-021 and sulbutiamine, and (4) Saline self-administration (control). On Day 15, subjects were sacrificed following the last session. A control group was examined as described above, but subjects received intravenous saline infusions, instead of fentanyl. Catheters were flushed daily with physiological sterile saline solution containing heparin. Catheter integrity was verified at the time of sacrifice.

#### Tissue Histology

Liver samples were collected and fixed in in 4% paraformaldehyde. Subjects were then perfused through the heart with RNAse-free PBS, decapitated, and brains were removed. The cerebellum from the brain was fixed in in 4% paraformaldehyde for histological analysis. For histological analyses, analysis was performed by Reveal Biosciences with a board-certified veterinary pathologist evaluation. Liver and brain tissues were frozen, embedded in paraffin and sectioned. Sections were collected from each sample at 4μm and stained with hematoxylin and eosin (H&E). Images were evaluated for biliary hyperplasia, portal leukocytes, cytoplasmic vacuoles, hepatocellular fatty change, and hepatic necrosis. Findings were scored as 0=within normal limits, 1=minimal or slight (least extent discernible), 2=mild, 3=moderate, 4=marked, and 5=severe.

#### Statistics

Data were analyzed by a one-way or two-way analysis of variance (ANOVA) using Graphpad Prism software, as appropriate. Significant main or interaction effects were followed by Dunnett or Sidak post-hoc comparison with correction for multiple comparisons. The criterion for significance was set at α=0.05.

## RESULTS

### In Vitro GATC-021 Analyses

In these studies, we characterized the effects of GATC-021 in mediating serotonergic signaling as a therapeutic approach for OUD. The GATC-021 compound was synthesized as previously described and then validated for its effectiveness in acting on various serotonin receptors in cell-based assays (**Table 1**). Of note, the greatest efficacy of GATC-021 was found for the 5HT2A and 5HT6 receptors, with lesser effects found at 5HT2B and 5HT7D.

**Table 1.** *In vitro* assays to determine GATC-021 activity on serotonergic and BDNF receptor signaling. Percent efficacy determined relative to control agonist ligand with 10 μM concentration.

| Functional Assay | Mean | SD | % Efficacy |
| --- | --- | --- | --- |
| 5-HTR2A | 109720.00 | 2375.9 | 45 |
| 5-HTR2B | 107.39 | 0 | -1.1 |
| 5-HTR6 | 23320.00 | 56.6 | 28.9 |
| 5-HTR7D | 3520.00 | 169.7 | 0.5 |
| TrkB | 3900.00 | 28.3 | -3.5 |

### Effects of GATC-021 on general behavior

To next investigate whether GATC-021 induces effects on general behavioral outcome measures, subjects were injected with GATC-021 and assessed in the open field test.

The open field test is a commonly used behavioral test to assess exploratory and minor anxiety-related behaviors. Subjects were examined across four parameters: (1) total distance travelled, which is an indication of general activity levels, (2) freezing episodes in which the rat remains motionless, often in a crouched or hunched position, which is often associated with fear or anxiety, (3) mobile time, which represents the period when the rat is actively moving and includes behaviors such as walking, running, exploring, or engaging in other forward movement activities, and (4) immobile time, which refers to the duration in which the rat remains motionless or shows minimal movement and can include freezing behavior or stationary periods. We examined a moderate (50 mg/kg) and higher (70 mg/kg) GATC-021 dose. For distance travelled, a significant difference was found among groups (*One-way Repeated Measures ANOVA,* F_(2, 10)_=5.941, p=0.0199) (**Figure 1A**). In the post-hoc analysis, it was determined that the higher 70 mg/kg GATC-021 dose decreased the overall distance travelled (p=0.0311), suggesting that this dose may exert general adverse effects. However, no statistically significant differences were found comparing 50 mg/kg dose to control, thereby demonstrating that this dose did not induce any deficits in general movement behaviors. For the other behavioral parameters, no differences were found with GATC-021 treatment with freezing episodes (*One-way Repeated Measures ANOVA,* F_(2, 10)_=0.5130, p=0.6136) (**Figure 1B**), mobile time (*One-way Repeated Measures ANOVA,* F_(2, 10)_=1.086, p=0.3744) (**Figure 1C**), or immobile time (*One-way Repeated Measures ANOVA,* F_(2, 10)_=1.086, p=0.3744) (**Figure 1D**). Given the effects found at the higher 70 mg/kg GATC-021 dose, we excluded this dose from the subsequent stages of the study and incorporated a lower dose of 25 mg/kg.

**Figure 1.**
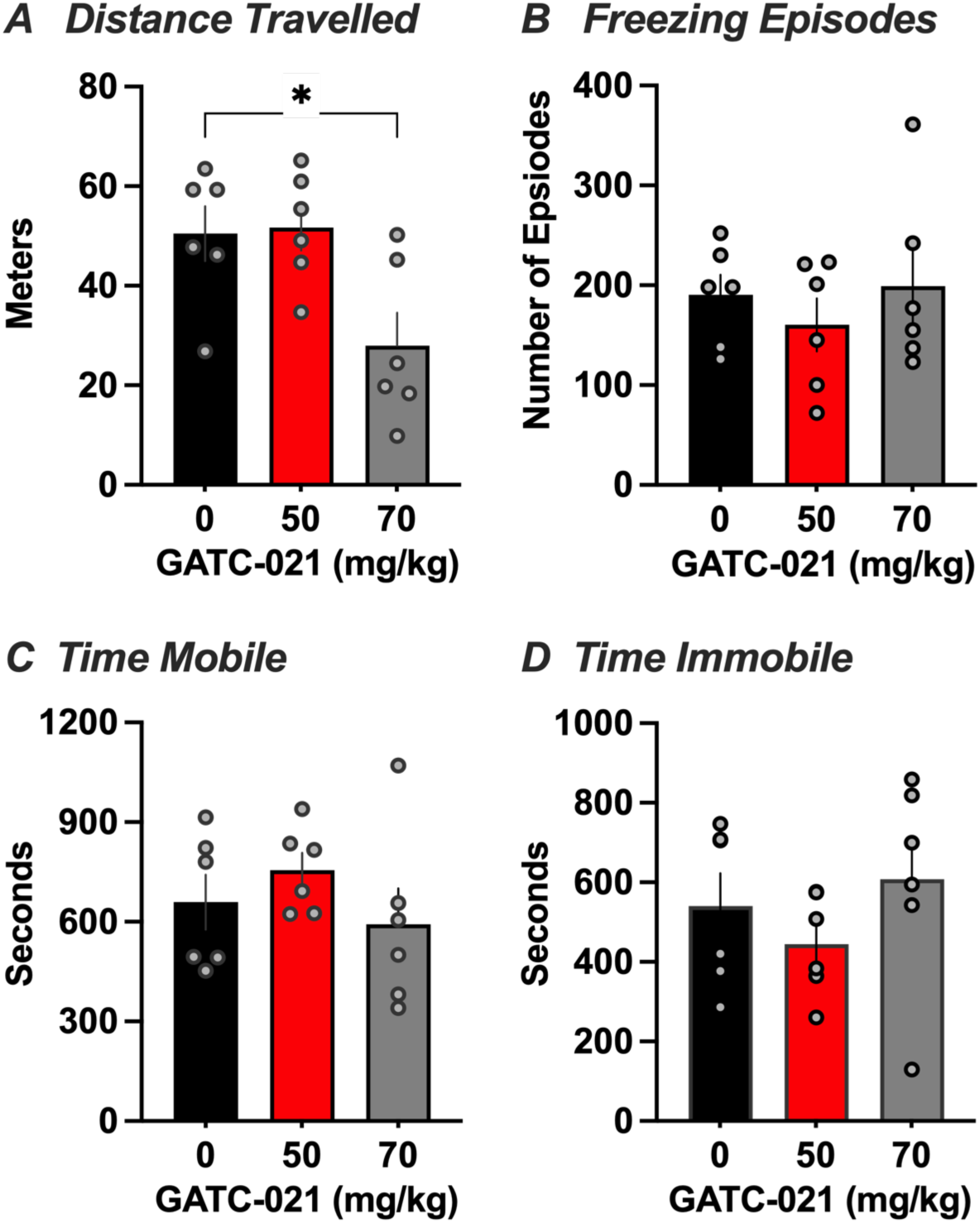
Effects of GATC-021 on general behavioral measures. Subjects were examined in the open field test with a within-subject design. (A) The highest 70 mg/kg dose of GATC-021 led to a reduction in overall distance travelled, but no differences were present comparing the lower 50 mg/kg dose relative to control. *p<0.05. (B-D) No differences among treatment groups were found in the total number of freezing episodes (B), time mobile (C), or time immobile (D). Data represent mean ± SEM, and individual subject values indicated with circles.

### Acquisition of Fentanyl Self-Administration

Male and female rats underwent intravenous fentanyl self-administration across 9 sessions at a dose of 2.5 μg/kg/infusion (*Two-way Repeated Measures ANOVA, Session* F_(8, 336)_=12.79, p<0.0001; *Sex* F_(1, 42)_=14.44, p=0.0005; *Interaction* F_(8, 336)_=1.626, p=0.1163) (**Figure 2A**). The post-hoc analysis revealed significantly greater intake by the female rats on session 1 (p=0.0004), 8 (p<0.0001) and 9 (p=0.0325). Both males and females demonstrated preference for the active lever associated with fentanyl infusions (*Two-way Repeated Measures ANOVA, Session* F_(8, 672)_ = 14.21, p<0.0001; *Sex/Lever* F_(3, 84)_=47.67, p<0.0001; *Interaction* F_(24, 672)_=10.52, p<0.0001) (**Figure 2B**). Comparing between active and inactive lever with the post-hoc, the male subjects demonstrated a significant preference for the active lever across sessions 5 (p=0.0004), 6 (p=0.0016), 7 (p<0.0001), 8 (p<0.0001), and 9 (p<0.0001). Female subjects learned the active versus inactive lever pressing behavior requirements quicker as they demonstrated a significant preference for the active lever across all sessions 1-9 (p<0.0001).

**Figure 2.**
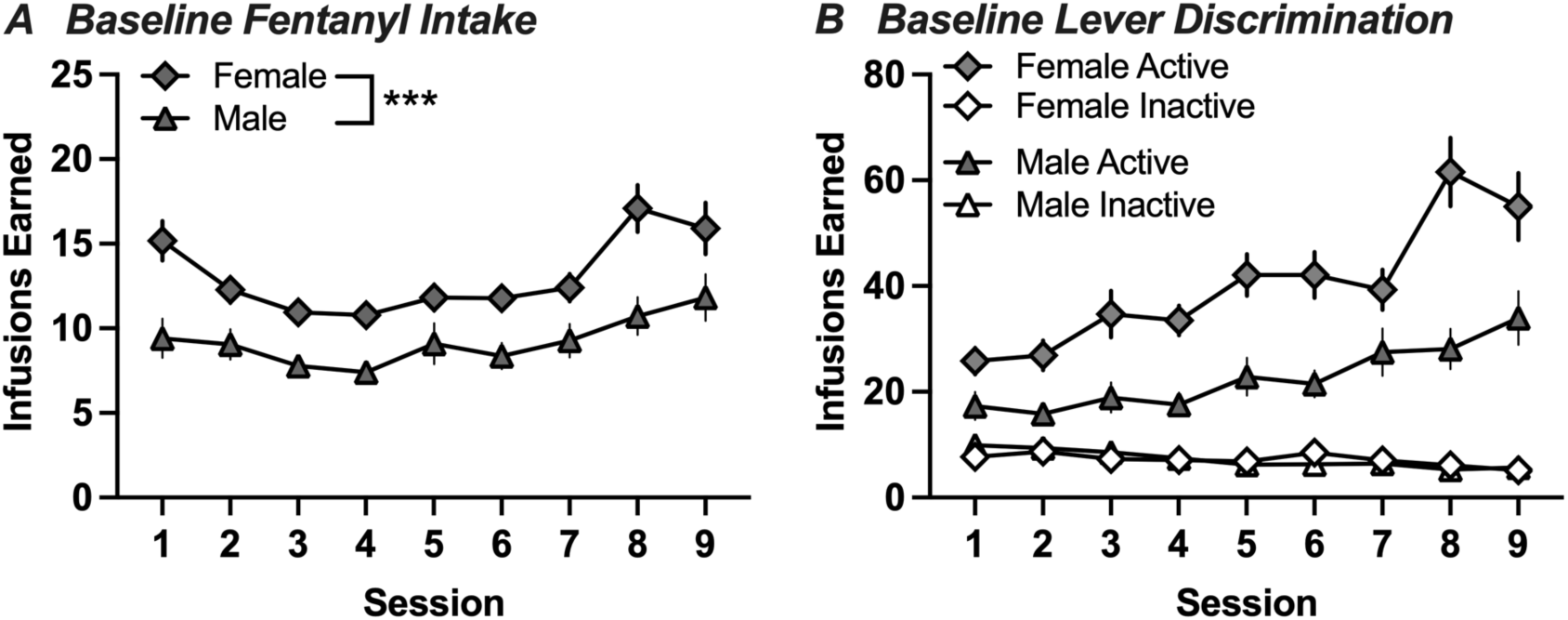
Intravenous fentanyl self-administration acquisition across sessions. (A) While both males and females demonstrated fentanyl intake, the females earned an overall higher number of drug infusions across the testing period. ***p<0.0001 (B) When comparing the active and inactive lever pressing behavior, both males and females exhibited a preference for the active lever, thereby demonstrating preferential lever-pressing to receive fentanyl infusions. Data represent mean ± SEM.

### Effects of GATC-021 on Fentanyl Intake

We next analyzed whether GATC-021 could alter fentanyl intake following an acute dose or extended daily dosing period. In male subjects, the acute injection of the GATC-021 compound exerted a reduction in overall fentanyl intake (*One-way Repeated Measures ANOVA,* F_(2, 16)_=5.571, p=0.0146) (**Figure 3A**). The post-hoc test revealed that a significant decrease in overall fentanyl intake was induced at both the 25 mg/kg (p=0.0481) and 50 mg/kg (p=0.0481) doses. After five days of GATC-021 dosing, significant differences were still present for the effect in reducing fentanyl intake, but only with the moderate 50 mg/kg dose (*One-way Repeated Measures ANOVA,* F_(2, 16)_= 3.474, p=0.0558; post-hoc test, 0 vs. 50 mg/kg, p=0.0360) (**Figure 3B**). These data demonstrate that GATC-021 had a significant impact in reducing fentanyl intake in male subjects, and the moderate dose partially maintained its effects across five consecutive days, indicating a potential for sustained effects with more long-term dosing.

**Figure 3.**
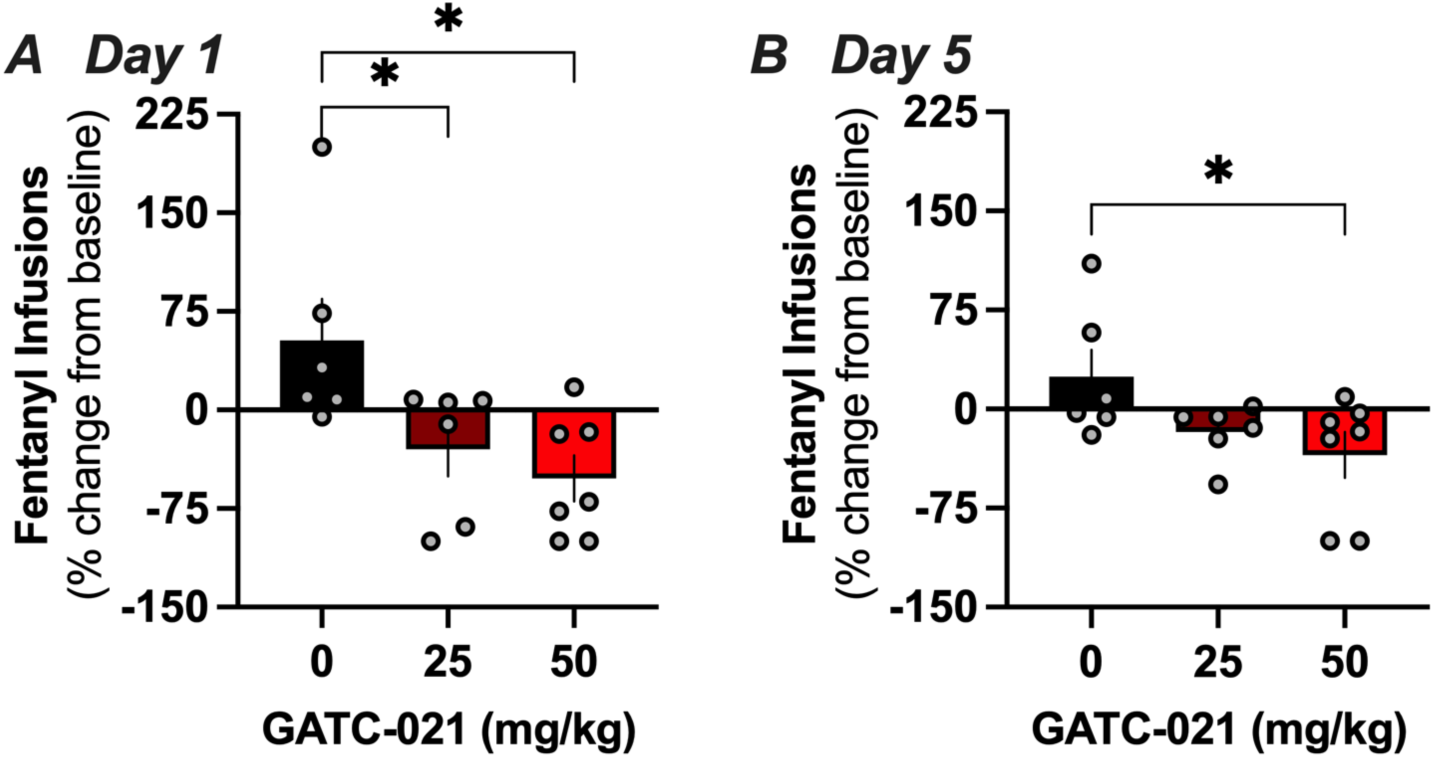
GATC-021 reduces fentanyl intake in males. (A) Administration of either the low or moderate dose of GATC-021 led to a reduction in the infusions earned on the first day of injections. *p<0.05 (B) On the fifth consecutive day of GATC-021 injections with fentanyl self-administration, the moderate dose retained some effectiveness in reducing fentanyl intake. Data represent mean ± SEM, and individual subject values indicated with circles. *p<0.05

Next, females were examined for the effects of GATC-021 on drug taking behavior. With acute administration, the moderate dose of 50 mg/kg exerted a significant reduction in fentanyl intake (*One-way Repeated Measures ANOVA,* F_(2, 15)_=6.390, p=0.0098; post-hoc test 0 vs. 50 mg/kg, p=0.0052) (**Figure 4A**). After five days of consecutive dosing, this significant attenuation of fentanyl was maintained (*One-way Repeated Measures ANOVA,* F_(2, 15)_=4.611, p=0.0275; post-hoc test 0 vs. 50 mg/kg, p=0.0169) (**Figure 4B**). These data demonstrate that the moderate dose of GATC-021 had a significant impact in reducing fentanyl intake in female subjects with both acute and longer-term dosing.

**Figure 4.**
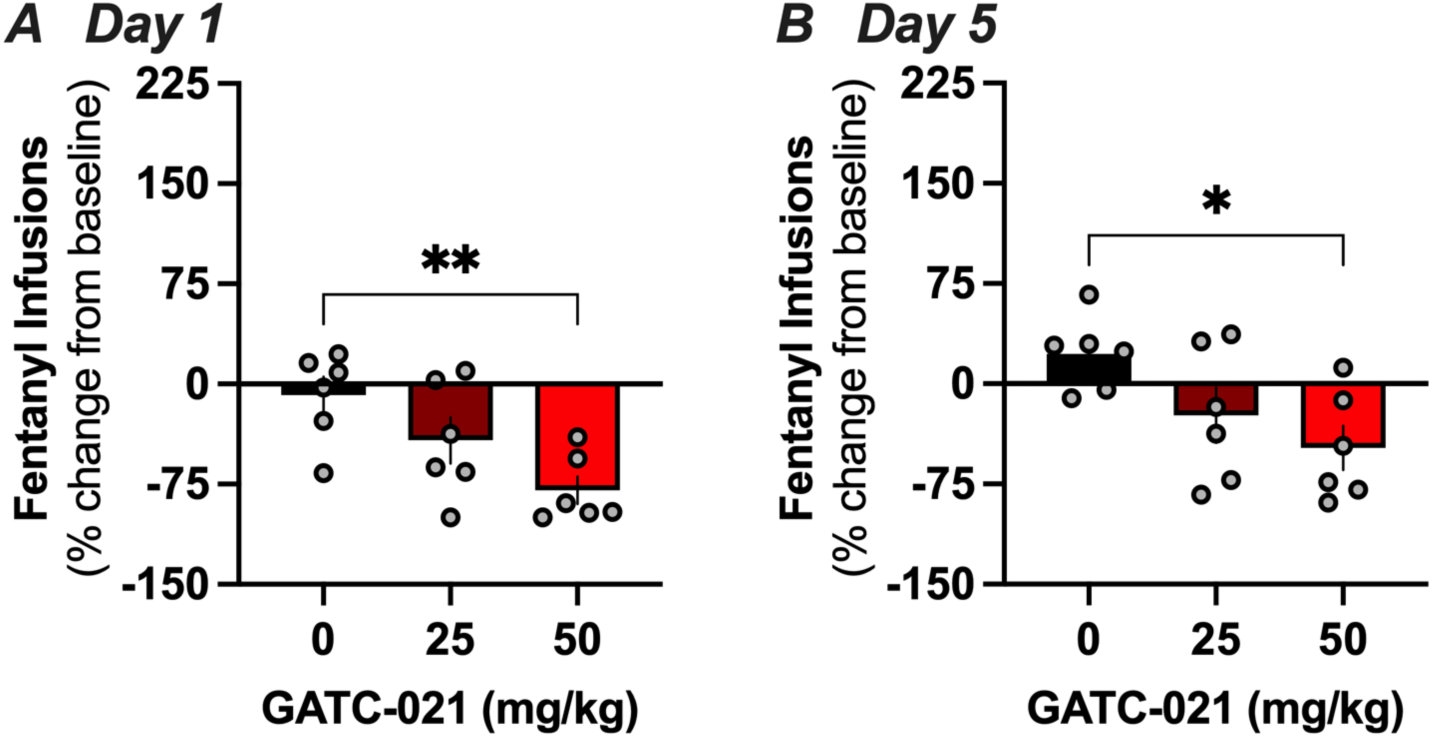
GATC-021 reduces fentanyl intake in females. (A) Administration of the moderate dose of GATC-021 led to a reduction in the infusions earned on the first day of injections. **p<0.01 (B) On the fifth consecutive day of GATC-021 injections with fentanyl self-administration, the moderate dose retained effectiveness in reducing fentanyl intake in females. Data represent mean±SEM, and individual subject values indicated with circles. *p<0.05

### No gross abnormalities with liver and brain histological analyses

Given that the drug formulation was observed to precipitate out of solution within a few minutes after deriving the pH, which necessitated immediate injections following solution preparation, concerns arose regarding potential organ toxicity due to intraperitoneal injection location. Thus, liver sections were analyzed for abnormal histological markers (*Two-way Repeated Measures ANOVA, Treatment* F_(3, 46)_=0.7392, p=0.5341; *Histological Indication* F_(5, 230)_=148.9, p<0.0001; *Interaction* F_(15, 230)_=1.114, p=0.3445) (**Figure 5**), and no significant differences were noted between groups in the post-hoc analysis. Overall, the liver tissue exhibited portal leukocytes, observed as infiltrates of leukocytes including macrophages, lymphocytes, and neutrophils, which were located in proximity to portal areas. Sinusoidal leukocyte foci were characteristic of granulomas, with the predominant cells being classified as macrophages. While overall similar among treatment groups, portal leukocytes and focal sinusoidal leukocytes were slightly more notable (minimal to moderate) in the untreated subjects, but no statistically significant differences were found between treatment groups. Hepatocytes with intracytoplasmic hepatocellular fatty vacuoles were classified as hepatocyte fatty change, which was similar across all treatment groups. Cytoplasmic vacuoles were fairly similar in all groups; cytoplasmic vacuoles can reflect feeding status, metabolism, or an artifact from fixation (e.g., freezing and thawing of tissue). Biliary hyperplasia was characterized by increased total number of bile ducts and/or clusters of epithelial cells in bile ducts and was minimal to mild in all groups. One liver section in the fentanyl group that received the 25 mg/kg GATC-021 dose had moderately large foci of necrosis with neutrophils, macrophages, lymphocytes, and necrotic hepatocytes, but this pathology was not noted in the other subjects receiving the same treatment. All brain sections were within the range of normal with no abnormal findings. Thus, these findings support the conclusion that GATC-021 did not induce adverse effects in organ systems.

**Figure 5.**
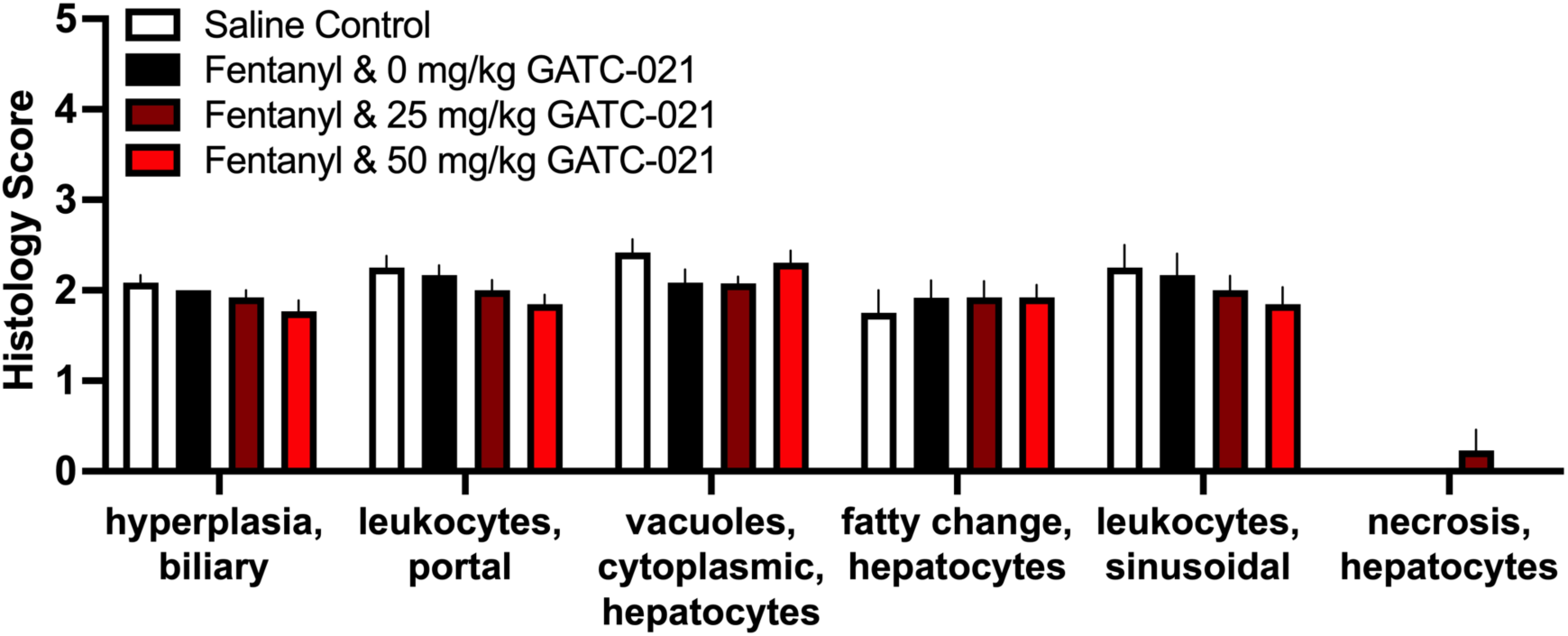
Normal histological findings for liver tissue. Groups did not differ across the histological indications. For each noted indication, sections were scored as 0=within normal limits, 1=minimal or slight (least extent discernible), 2=mild, 3=moderate, 4=marked, and 5=severe. Data represent mean ± SEM.

## DISCUSSION

In these studies, we found that GATC-021 combined with sulbutiamine reduced fentanyl intake in both male and female rats, with the 50 mg/kg dose being most efficacious. These effects on drug intake were selective in reducing fentanyl intake, as no differences were found at this dose in control behavioral measures. Moreover, no histological indications of potential adverse effects with liver or brain tissue were observed based on group treatment condition. Given that GATC-021 exerts partial agonist actions at the 5HT2A and 5HT6 receptors, these findings suggests that modulation of these targets are a viable approach for therapeutic development to promote the cessation of opioid use. Further work is underway to understand the role and ratio of targeting, as well as the role of thiamine.

### Considerations for Sex Differences

At baseline, females exhibited a higher level of fentanyl intake than males. These findings are in accordance with prior reports from the field, which further suggest that these differences are likely regulated by estradiol levels^24,25^. Given this higher level of drug consumption, it was important to examine the effects of GATC-021 in both male and female subjects across doses. Overall, the GATC-021 compound attenuated fentanyl intake in a similar manner in both males and females. However, it is worth noting that the specific effects of each dose appeared to differ between sexes, but importantly, the moderate dose of 50 mg/kg GATC-021 showing a significant impact in both sexes.

### Acute Versus Longer-Term Dosing Paradigms

Differential effectiveness was found based on the duration of dosing across fentanyl self-administration days. While the moderate dose of 50 mg/kg maintained its effectiveness on day 5, the lower dose of 25 mg/kg lost its significant effects in the male subjects. This suggests that the effects of the GATC-021 compound may vary over time, thus emphasizing the importance of considering the duration of treatment for its impact on drug intake to support translational potential in the clinical setting. This also raises the possibility of effectiveness in attenuating relapse behaviors following a period of abstinence, which will need to be investigated in the future.

### Conclusions

Our findings provide validation that non-hallucinogenic 5-HT receptor targeting reduces indications of OUD in preclinical models. Further research is warranted to better understand the pharmacokinetics associated with the GATC-021 compound, to explore potential strategies for optimizing dosing given solubility issues, and to understand the impact of GATC-021 on underlying neural substrates. Taken together, these novel findings serve as a foundation for further clinical development to provide a means for individuals to overcome opioid addiction and end the opioid epidemic.

## Acknowledgements and Disclosures

This project was supported by a sponsored research project from GATC Health. We would like to thank the West Virginia University Shared Research Facilities for access to HRMS instrumentation. SM, ACM, IJ, JRTL, and RT are GATC Health employees, and JRTL and CDF serve on the scientific advisory board for GATC Health.

## Author Contributions

The initial conceptualization of the project was developed by IJ, RT and JRTL. Chemistry synthesis of the compound was performed by RT, and structure validation was performed by ACM. Overall project administration was overseen by SM and IJ, including data curation for the *in vitro* screens and histological tissue analyses. Software for the AI platform was developed by IJ. *In vivo* studies were conceptualized, methodology derived, data analyzed and validated, and statistical analyses performed by VL and CDF. *In vivo* experiment investigation and tissue collection was performed by VL, JPF, MB, AM, JEC and CDF. The original draft was written by VL, SM, ACM, RT and CDF. The manuscript was revised by all authors, and the final version was approved by all authors.

## Supplementary Material

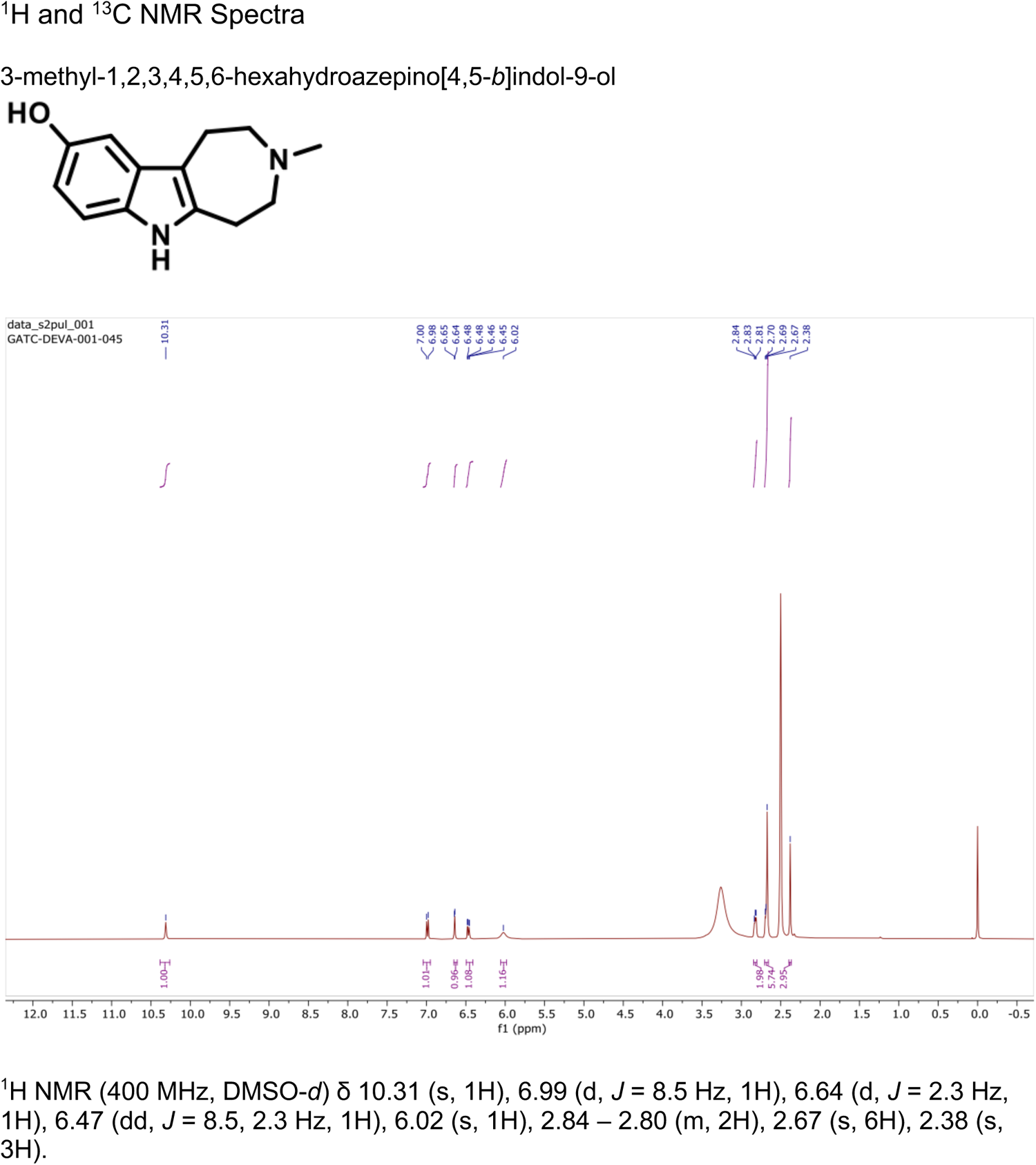
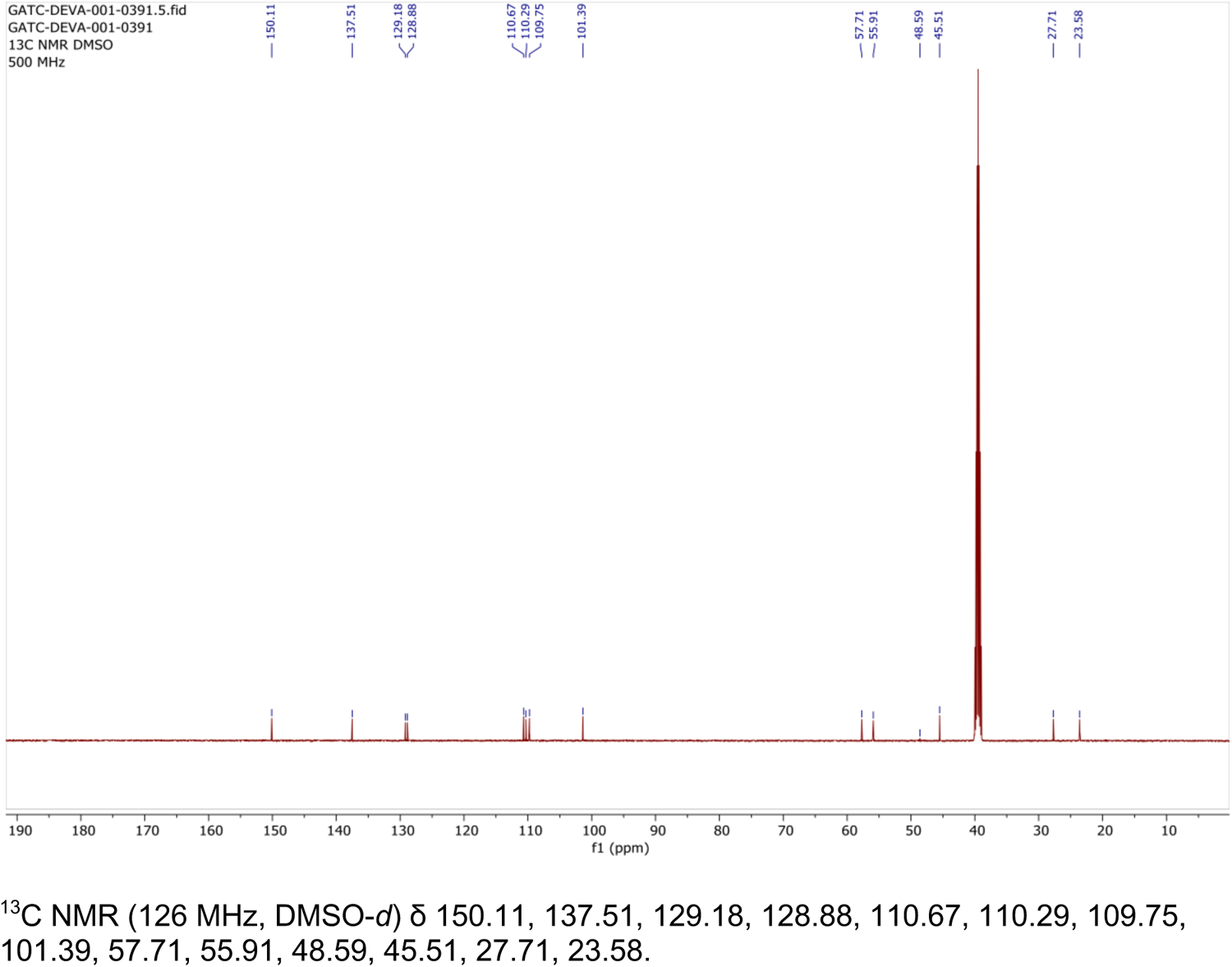

